# A xeno-free protocol for rapid differentiation of human iPSC-derived microglia from the KOLF2.1J reference line

**DOI:** 10.1101/2025.07.28.667242

**Authors:** Nélio A.J. Oliveira, Katherine R. Lewkowicz, Patricia A. Clow, Michael E. Ward, Mark R. Cookson, William C. Skarnes, Justin A. McDonough

## Abstract

We present a detailed, xeno-free protocol for the rapid differentiation of human induced pluripotent stem cells (hiPSCs) into microglia using the well-characterized KOLF2.1J reference line. This system employs doxycycline-inducible expression of six transcription factors (6-TF), stably integrated into the CLYBL safe harbor locus, to drive uniform microglial differentiation within two weeks. Adapted from Dräger et al. (1), our protocol includes key optimizations for KOLF2.1J, including culture on Laminin-521 to support xeno-free conditions. The resulting i-Microglia exhibit hallmark features of mature microglia, including expression of P2RY12, loss of the pluripotency marker SSEA4, phagocytic activity, and upregulation of immune markers (e.g., CD80, CD83) upon LPS stimulation. We also demonstrate compatibility with co-culture systems using iPSC-derived neurons. Additionally, we describe a modification of the line to include a constitutive mCherry reporter integrated into the SH4-2 safe harbor locus, enabling fluorescent tracking of microglia in mixed cultures or in vivo. This protocol provides a reproducible and scalable platform for generating functional human microglia from a widely used hiPSC line, supporting applications in disease modeling, neuroinflammation research, and therapeutic screening.

## Introduction

Microglia, the resident immune cells of the central nervous system, play critical roles in brain development, homeostasis, neurodegenerative diseases, and aging (2-5). Traditionally, microglia used in research are derived from animal models (primarily mice) or isolated from human tissue (6, 7). However, these sources present significant limitations due to interspecies differences and the limited availability and variability of primary human microglia.

Induced pluripotent stem cells (iPSCs) offer a renewable and scalable source for generating microglia and other cell types for basic and translational research (3, 8-10). Despite this, conventional iPSC differentiation protocols are often time-consuming, costly, and reliant on complex media and animal-derived components, which can introduce variability and limit clinical relevance (10, 11). Moreover, many existing iPSC-derived microglia models exhibit limited functional maturity, including poor responsiveness to immune stimuli and reduced phagocytic capacity, which constrains their utility in disease modeling and drug screening (3).

To address these challenges, transcription factor (TF)-based differentiation systems have been developed, enabling rapid and directed cell fate specification through forced expression of lineage-defining TFs. These TFs are typically stably integrated into the iPSC genome using transposon or knock-in strategies (1, 12, 13). A recent study□demonstrated□that human iPSC-derived microglia (i-Microglia) can be generated by inducible expression of six TFs– CEBPA, CEBPB, IRF5, IRF8, PU.1/SPI1, MAFB – resulting in cells with microglial morphology, phagocytic activity, and transcriptional profiles similar to primary microglia (1). However, further validation is needed to determine whether these cells can serve as reliable models for immune activation, drug screening, and integration into more complex neural systems.

In parallel, co-culture systems that incorporate both neurons and microglia are essential for modeling cell-cell interactions relevant to neurodegenerative and neuroinflammatory diseases, yet remain underdeveloped in scalable, defined formats. Additionally, microglia transplantation is emerging as a potential therapy for neuroinflammatory conditions such as Alzheimer’s disease, Parkinson’s disease, and central nervous system injuries (14, 15). For translational applications such as cell therapy or high-throughput screening, xeno-free protocols are preferred to avoid risks associated with animal-derived components, including immunogenicity and non-human post-translational modifications (16, 17).

In this study, we developed a rapid, xeno-free, transcription factor-driven protocol for generating functional human iPSC-derived microglia using the well-characterized reference iPSC line KOLF2.1J (18). We demonstrate that these cells exhibit key microglial functions, including phagocytosis and immune activation, and can be stably co-cultured with iPSC-derived cortical glutamatergic neurons. Our system provides a robust and versatile platform for studying human microglia biology, modeling disease, and testing therapeutic interventions.

## Results

### Generation of inducible i-Microglia cell lines

The inducible microglia (i-Microglia) line was engineered in KOLF2.1J cells (18) using CRISPR/Cas9-mediated insertion of six TFs (1) into the CLYBL safe harbor locus. The six TFs were divided across both CLYBL alleles via knock-in of two separate constructs. Each construct contained three TFs separated by T2A elements and driven by a TetON inducible promoter: PU.1/SPI1-CEBPB-IRF5 and MAFB-IRF8-CEBPA (Figure 1). Both constructs also included the reverse tetracycline transactivator (rtTA) under a constitutive promoter, enabling doxycycline-dependent co-expression of all six TFs (19).

**Figure 1.**
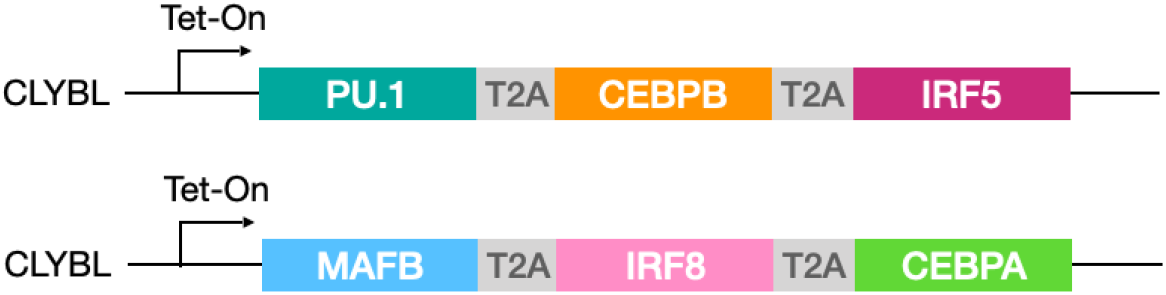
Six transcription factors (PU.1/SPI1, CEBPB, IRF5, MAFB, IRF8, CEBPA) were inserted into both CLYBL alleles of KOLF2.1J iPSCs using CRISPR/Cas9. Each allele received a Tet-ON-inducible construct encoding three TFs separated by T2A elements. A constitutive rtTA enables doxycycline-dependent co-expression of all six TFs.

We selected two engineered clones, A02 and D01, for further characterization. Both clones exhibited a normal karyotype and retained expression of pluripotency markers, as confirmed by immunofluorescence (IF) for OCT4 and SSEA4, and by flow cytometry (FC) for SSEA4 (Supplemental Figure S1). As expected, FC analysis showed that undifferentiated iPSCs were negative for P2RY12, a microglia-specific marker, indicating that the inserted constructs remained transcriptionally silent in the absence of doxycycline and did not exhibit significant leaky expression.

To create a fluorescent reporter version of the i-Microglia line suitable for both *in vitro* and *in vivo* applications, we introduced a constitutively expressed mCherry transgene into a newly identified safe harbor locus on chromosome 4, designated SH4-2 (20), in i-Microglia clone A02. Following transgene integration, we isolated two clones, F04 and G09, both of which exhibited near 100% mCherry expression (Supplemental Fig S2). Importantly, mCherry expression remained stable after at least two additional passages, confirming that the SH4-2 locus supports sustained transgene expression at the iPSC stage.

### Optimization of substrate conditions for iPSC-to-microglia differentiation

To establish a robust and reproducible differentiation protocol for our i-Microglia line, we first evaluated the previously published protocol for the 6-TF system (1). Initial differentiation of iPSC clones (A02 and D01) was performed on dual-coated surfaces—Poly-D-Lysine (PDL) and Matrigel—as described in the original protocol with minor modifications. Doxycycline (DOX) was added at day 0 to induce TF expression and withdrawn on day 8 (Figure 2A). We also supplemented the culture with CX3CL1 on day 8, a chemokine known to support microglial homeostasis. By day 6, cells exhibited microglia-like morphology and expressed canonical markers such as IBA1 and CD68 by days 10–15 (Supplemental Figure S3), confirming compatibility of the system with our cell line.

**Figure 2.**
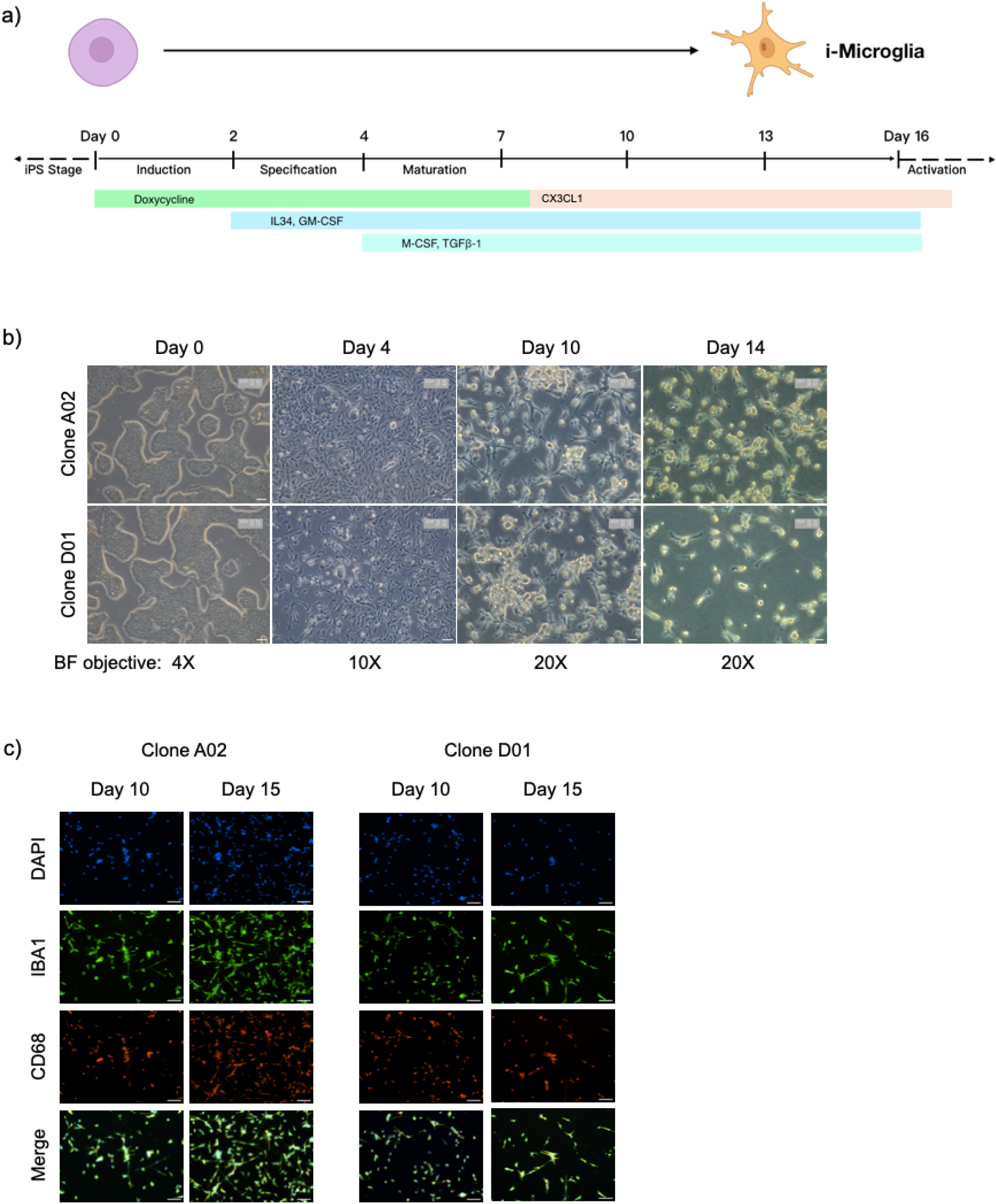
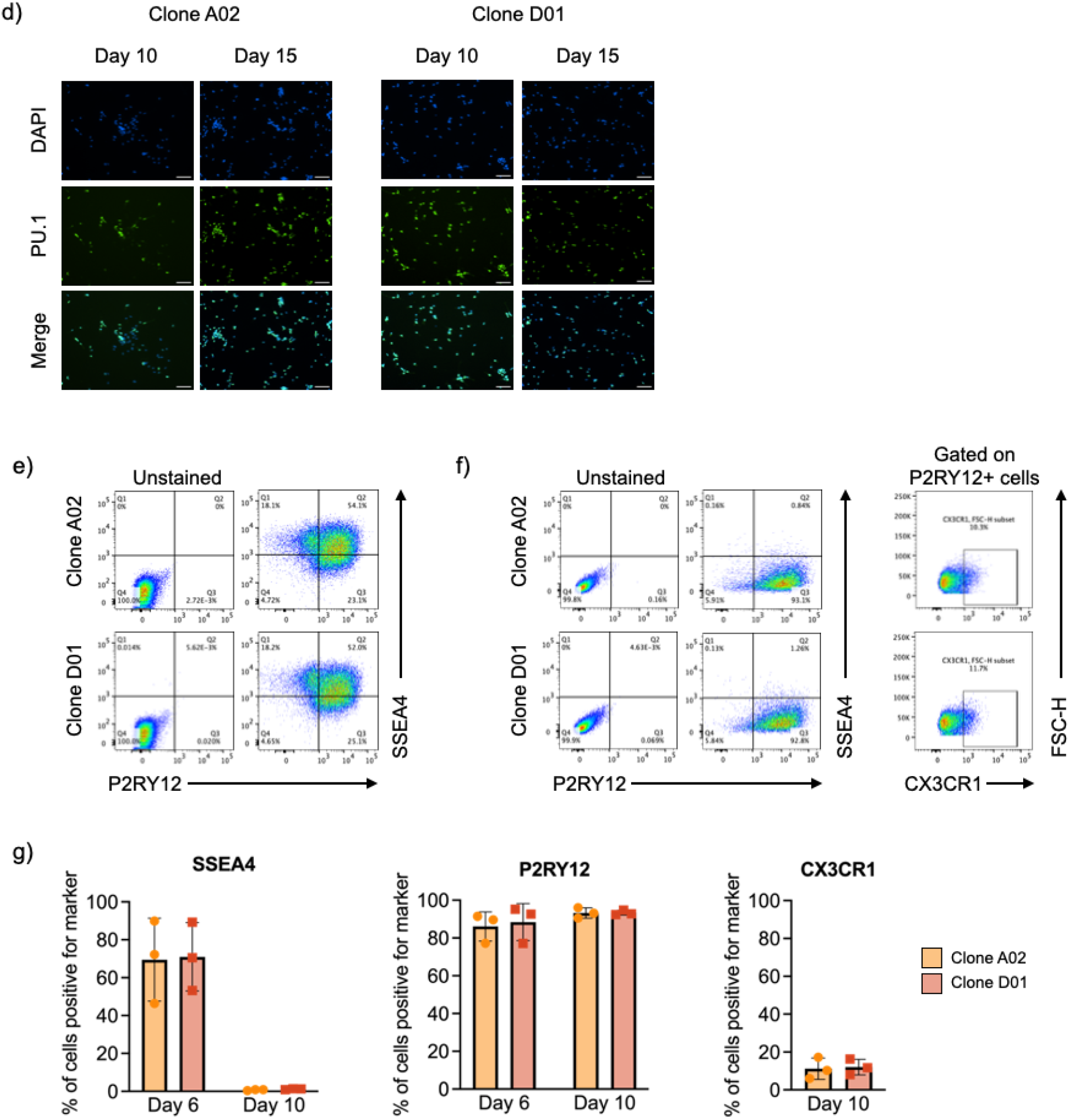
Differentiation of i-Microglia on Poly-D-Lysine + Laminin-521 Substrate. a) Schematic timeline of the 15-day i-Microglia differentiation protocol. From day 0 to day 2 (induction stage), iPSCs were cultured in mTeSR+ medium with 2□µg/mL doxycycline to induce expression of six transcription factors. From day 2 to day 4 (specification stage), cells were cultured in Advanced DMEM/F12 supplemented with 100□ng/mL IL-34 and 10□ng/mL GM-CSF. From day 4 onward (maturation stage), the medium was supplemented with 50□ng/mL M-CSF and 50□ng/mL TGF-β1. On day 8, CX3CL1 was added and doxycycline was withdrawn. Media was refreshed every 2–3 days thereafter. b) Brightfield images showing morphological progression of KOLF2.1J iPSCs (clones A02 and D01) from day 0 (undifferentiated) to days 4, 10, and 14. c) Immunofluorescence (10× objective) at days 10 and 15 showing expression of microglial markers IBA1 (green) and CD68 (red), with DAPI nuclear stain (blue), in clones A02 and D01. d) Immunofluorescence (10× objective) of a separate well showing PU.1 (green) and DAPI (blue) at days 10 and 15. e) Representative flow cytometry plots at day 6 showing P2RY12 (x-axis) and SSEA4 (y-axis) expression in clones A02 and D01, comparing stained versus unstained controls. f) Flow cytometry plots at day 10 using the same staining conditions as in (e), along with additional plots showing CX3CR1 (x-axis) versus FSC-axis) gated on P2RY12-positive cells. g) Quantification of flow cytometry data from three biological replicates (y-showing the percentage of cells positive for SSEA4 (left), P2RY12 (middle), and CX3CR1 (right) in clones A02 and D01. Error bars represent mean ± SEM.

We next sought to transition toward a xeno-free protocol and therefore tested human recombinant laminins as alternatives to Matrigel. Cells were plated on PDL with either Laminin-111 (LAM111) or Laminin-521 (LAM521), and differentiation was initiated with DOX on day 0. Both laminin conditions supported early microglial morphology and expression of key markers (IBA1, CD68, PU.1, and P2RY12) by day 6 (Supplemental Figure S4). However, only LAM521 consistently supported cell viability and marker expression through the full 15-day differentiation period (Figure 2B–G), whereas LAM111 showed reduced cell survival (Supplemental Figure S4).

Based on these results, we selected PDL + LAM521 as the optimized, xeno-free substrate for all subsequent experiments. This condition supported robust expression of microglial markers (IBA1, CD68, PU.1, P2RY12, CX3CR1) and complete loss of pluripotency marker SSEA4 over time (Figure 2C–G), confirming successful and reproducible differentiation. Notably, PU.1 expression persisted for 7 days after DOX withdrawal (day 15), indicating that its sustained expression reflects endogenous activation in mature microglia rather than continued transgene induction (Figure 2D). P2RY12 was detectable as early as day 6 under all conditions, with expression peaking at over 90% of cells by day 10 (Figures 2E-G).

Similarly, i-Microglia-mCherry clone F04 maintained stable mCherry expression throughout differentiation under the optimized xeno-free conditions, and mCherry-positive microglia remained positive for P2RY12 (Figure 3). These results confirm that the SH4-2 safe harbor locus supports stable transgene expression during differentiation.

**Figure 3.**
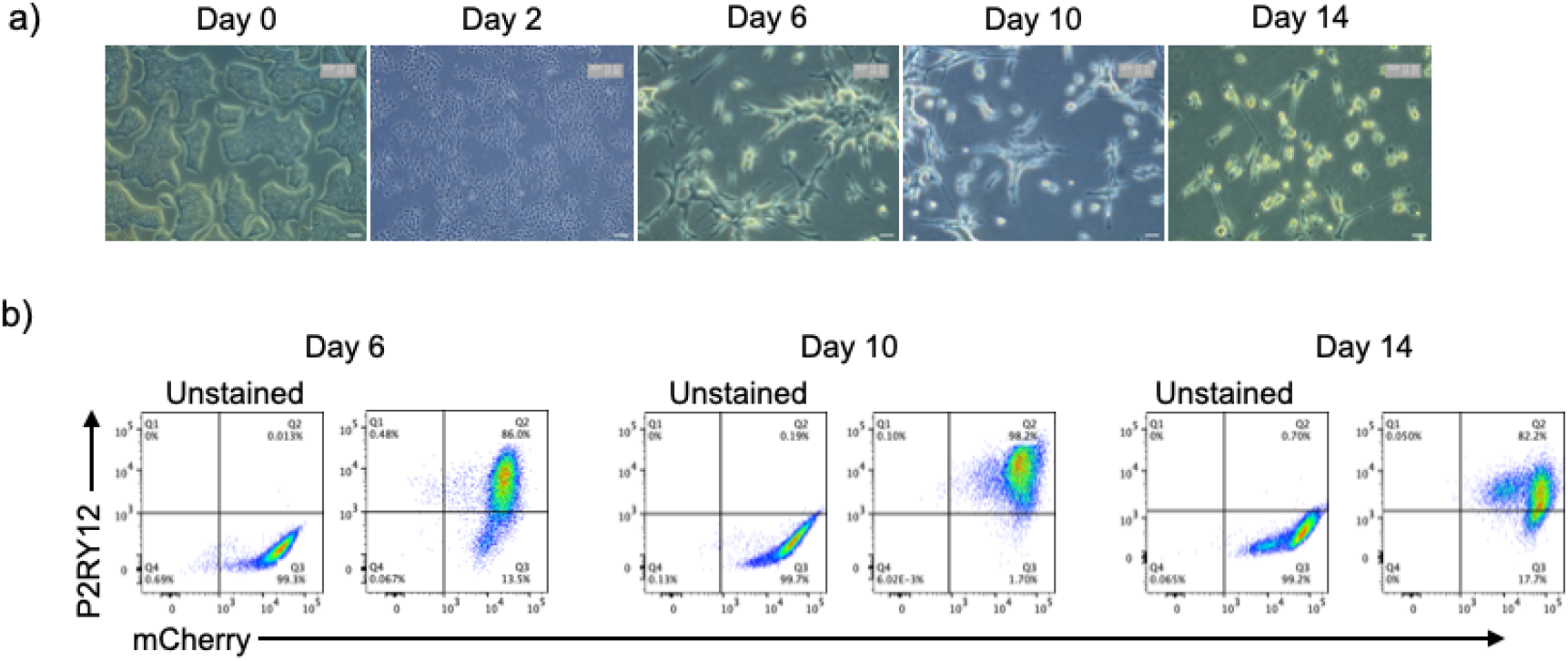
Time-Course Differentiation and Characterization of i-Microglia-mCherry. a) Brightfield images showing morphological progression of clone F04 during differentiation. Images were acquired using a 4x objective at the day 0 iPSC stage and day 2, and a 20x objective at days 6, 10, and 14. b) Flow cytometry analysis showing stable mCherry expression (x-axis) and P2RY12 expression (y-axis) at days 6, 10, and 14 under optimized Poly-D-Lysine + Laminin-521 coating conditions.

### Phagocytosis

To assess the functional capacity of i-Microglia differentiated on Poly-D-Lysine and LAM521, we performed a phagocytosis assay using zymosan beads labeled with a pH-sensitive green-fluorescent dye (pHrodo). By day 9 of differentiation, most cells exhibited robust phagocytic activity. Flow cytometry analysis confirmed that over 75% of the cell population internalized the labeled beads within 2.5 hours (Figure 4A). For confocal imaging, we used the i-Microglia-mCherry clone F04, differentiated for 10 days. Only cells that successfully phagocytosed the zymosan particles exhibited green fluorescence (Figure 4B). Live-cell imaging further demonstrated the dynamic nature of phagocytosis in both parental clones (A02 and D01) and the mCherry-expressing clone F04. In all cases, cells turned green upon internalization and acidification of the zymosan particles (Supplemental Videos 1 and 2), providing real-time evidence of functional microglial behavior.

**Figure 4.**
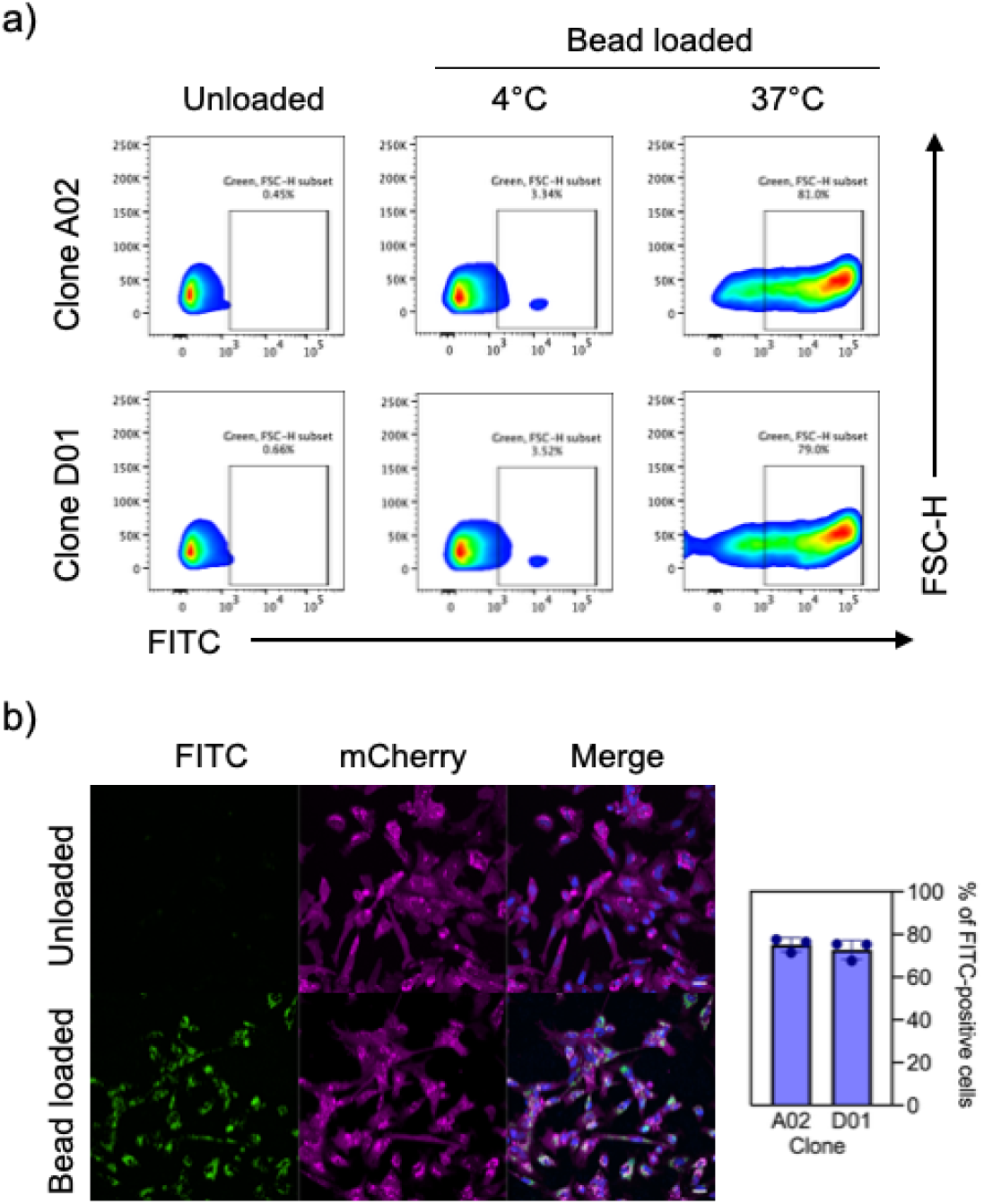
Phagocytic Activity of i-Microglia. a) Representative flow cytometry plots showing uptake of green fluorescent (FITC) zymosan beads by i-Microglia clones A02 and D01. Cells were incubated with beads at 37°C to assess active phagocytosis or at 4°C as a binding-only control. FITC fluorescence is shown on the x-axis. b) Confocal microscopy images of mCherry-labeled i-Microglia clone F04 incubated with or without FITC-labeled zymosan beads. Images were acquired using a 40x objective.

### Upregulation of i-Microglia activation markers upon stimulation

A common limitation of *in vitro* iPSC-derived microglia models is their incomplete maturation and limited responsiveness to immune stimuli (3). To assess the immunocompetence of our i-Microglia, we evaluated the expression of key immune activation markers under both basal and lipopolysaccharide (LPS)-stimulated conditions. i-Microglia differentiated for at least 10 days were treated with 0.5□µg/mL LPS for 22 hours, while media-only controls were included for comparison. Flow cytometry analysis revealed a modest increase in P2RY12 expression following LPS stimulation in both clones A02 and D01 (Figure 5A), suggesting a potential shift in microglial activation state.

**Figure 5.**
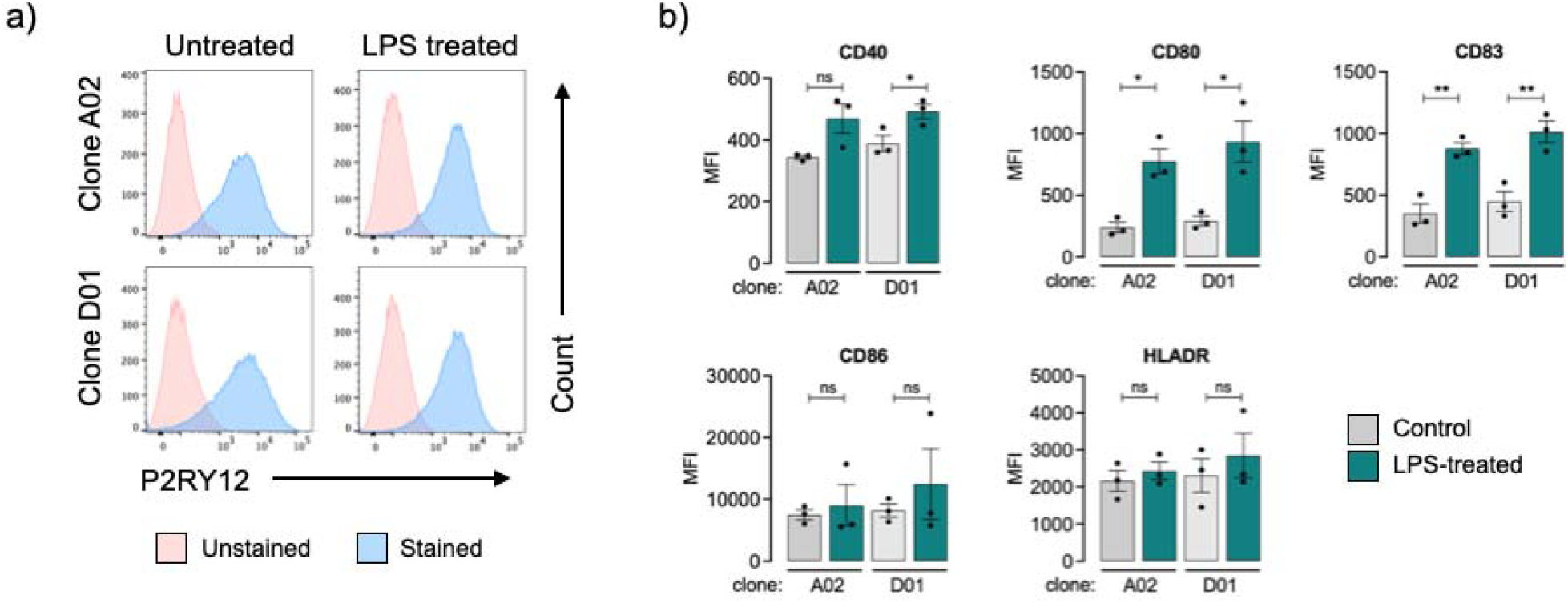
Immune Activation of i-Microglia Following LPS Stimulation. a) Representative flow cytometry plots showing P2RY12 expression in clones A02 and D01 under control (untreated) and LPS-treated conditions. Unstained and stained populations are shown for each condition. b) Quantification of immune activation markers in P2RY12-positive cells following LPS stimulation. Bar graphs show mean fluorescence intensity (MFI) for CD40, CD80, CD83, CD86, and HLA-DR in clones A02 and D01 under control and LPS-treated conditions. Data represent mean ± SEM from three biological replicates. ^*^ = p<0.05, ^**^ = p<0.01, ns = not significant.

To further characterize the immune response, we gated on the P2RY12-positive population and measured the expression of five activation markers: CD40, CD80, CD83, CD86, and HLA-DR (Figure 5B). All markers were detectable at baseline, indicating a degree of functional maturity by day 11. Upon LPS stimulation, all five markers were upregulated to varying extents in both clones. Notably, CD80 and CD83 showed statistically significant increases in both A02 and D01. CD40 was significantly upregulated in D01 but not A02, while CD86 and HLA-DR showed elevated expression in both clones without reaching statistical significance. These results demonstrate that KOLF2.1J-derived i-Microglia can mount a robust innate immune response upon stimulation.

### Establishing a co-culture system using inducible cortical glutaminergic neuron and i-Microglia

*In vitro* models that recapitulate the cellular complexity of brain tissue are essential for studying neurodegenerative and autoimmune diseases and for developing new therapeutic strategies. To evaluate the compatibility of our i-Microglia with neurons, we established a co-culture model using iPSC-derived cortical glutamatergic neurons (i-Neurons) generated via DOX-inducible expression of the TF neurogenin 2 (NGN2), as described by Flores et al. (21). i-Neurons were differentiated for 11 days following the established protocol. On day 4 of i-Microglia differentiation on Poly-D-Lysine and LAM521, cells were dissociated and seeded into the i-Neuron cultures at a 1:1 cell ratio. Each well of a 24-well plate received an equal number of i-Microglia and i-Neurons. To promote microglia attachment, either LAM521 or Laminin-3D (R&D Systems) was added to the i-Microglia medium. These co-cultures were maintained for at least ten days and subsequently stained for beta-III tubulin (TUJ1) to label neurons and IBA1 to label microglia. i-Microglia successfully integrated into the neuronal cultures under both attachment conditions (Figure 6), with clear expression of TUJ1 and IBA1 confirming the presence of both cell types. This co-culture system demonstrates the adaptability of i-Microglia to complex environments and provides a foundation for more advanced studies, such as modeling neuroinflammation, synaptic pruning, or neuron-microglia interactions in disease-relevant contexts.

**Figure 6.**
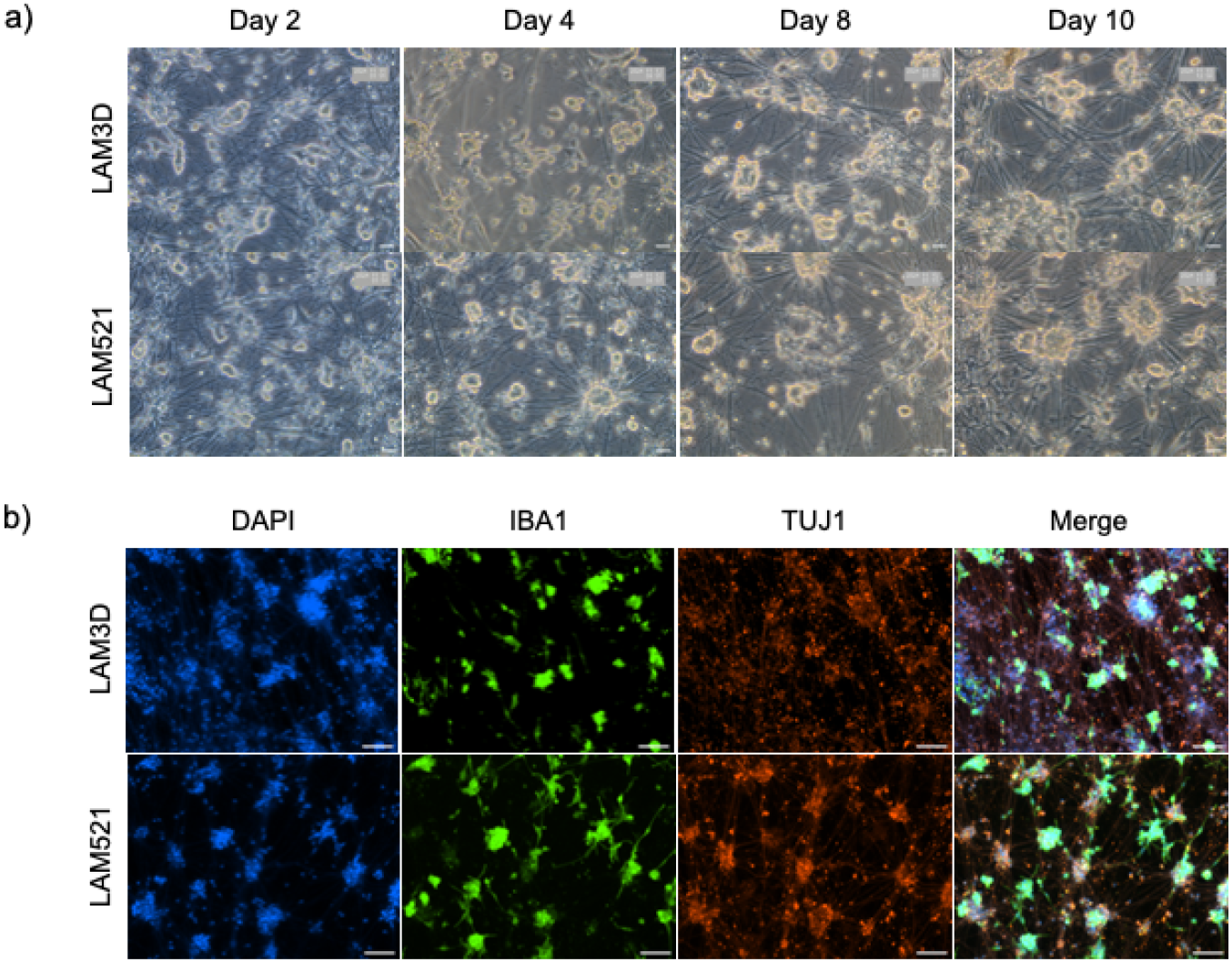
Co-Culture of Cortical Glutamatergic Neurons and i-Microglia. a) Brightfield time-course images of co-cultures at days 2, 4, 8, and 10, showing progressive neuronal network formation and microglial integration. b) Immunofluorescence at day 10 of co-culture showing TUJ1 (red, neuronal marker), IBA1 (green, microglial marker), and DAPI (blue, nuclei). Images acquired at 20× magnification for brightfield and 10× for immunofluorescence.

## Discussion

This protocol builds on the inducible microglia differentiation system developed by Dräger et al. (1), adapting it for use in the well-characterized human iPSC reference line KOLF2.1J (18). While the original 6-transcription factor (6-TF) system demonstrated rapid and efficient microglial induction, it had not been validated in KOLF2.1J, a line widely used for disease modeling and functional genomics. Our results confirm that the system performs robustly in this background, producing microglia with appropriate morphology, marker expression, and functional properties.

To enhance the stability and reproducibility of transgene expression, the 6-TF cassette was integrated into the CLYBL safe harbor locus, which has been shown to support higher and more consistent expression than commonly used sites such as AAVS1 (22, 23). AAVS1 is known to exhibit variable expression and silencing in myeloid cells, including microglia, while other loci such as CCR5 and Rosa26 present additional limitations, including potential susceptibility to viral infection or insufficient validation in human systems (24-27).

We also introduced two key modifications to the original Dräger et al. (1) protocol. First, we withdrew doxycycline (DOX) at day 8 rather than maintaining it throughout differentiation (M. Kampmann, personal communication). Our data show that this early withdrawal does not impair microglial identity or function. PU.1 expression persisted through day 15, suggesting that the microglial transcriptional program becomes self-sustaining after initial induction. This modification may better support natural transcriptional regulation. We speculate that prolonged DOX exposure could interfere with endogenous silencing mechanisms or alter microglial homeostasis, though further investigation is needed.

Second, we added CX3CL1 at day 8 (L. Erlebach, D. Kronenberg-Versteeg, personal communication). CX3CL1 is a chemokine commonly used in microglia differentiation protocols to support homeostasis and promote a more anti-inflammatory phenotype (3, 28, 29). Its inclusion likely contributes to the functional maturity and stability of the resulting i-Microglia.

A key advancement of our work is the successful generation of functional i-Microglia under xeno-free conditions, which enhances reproducibility and improves compatibility with translational and high-throughput applications. By eliminating animal-derived substrates such as Matrigel, our protocol avoids issues related to immunogenicity and non-human post-translational modifications, aligning with the growing preference for xeno-free systems in clinical and screening contexts (16, 17).

While the original protocol used Matrigel, we tested human recombinant laminins as alternatives to eliminate animal-derived components. Although Laminin-111 (LAM111) supported early differentiation, it failed to maintain cell viability through day 15. In contrast, LAM521 consistently supported robust differentiation and yielded over 90% P2RY12-positive cells. Progenitor cells exhibit lower affinity for LAM111 compared to LAM521 (30-34), which may explain its limited performance in our system. LAM521 has been widely used to support iPSC maintenance and differentiation into various lineages (35-37), and its role in supporting hematopoietic and myeloid cells is well documented (30, 32, 33).

Functionally, our i-Microglia demonstrated hallmark immune behaviors, including phagocytic activity and responsiveness to pathogen-associated molecular patterns (PAMPs). Stimulation with LPS led to significant upregulation of immune activation markers such as CD80 and CD83, indicating that our i-Microglia can mount a robust innate immune response. CD80 is a co-stimulatory molecule involved in antigen presentation and is typically expressed at low levels in homeostatic microglia (38, 39). Its upregulation upon LPS stimulation suggests that our cells can initiate adaptive immune responses (40, 41). CD83, which plays dual roles in promoting and resolving inflammation, was also significantly upregulated, further supporting the functional maturity of our i-Microglia (42-44).

To enhance the genetic tractability of our system, we validated a newly identified genomic safe harbor (GSH) locus, SH4-2, located in an intergenic region on chromosome 4 (20). Using an mCherry expression cassette containing a CAG promoter Woodchuck hepatitis virus-derived WPRE elements, we observed stable fluorescence throughout the entire microglia differentiation process. This finding is important, as there is growing demand for GSHs that support sustained transgene expression during differentiation, particularly in myeloid cells. mCherry-labeled i-Microglia are well suited for both in vitro and in vivo applications, including co-culture and potential transplantation studies.

Finally, we demonstrated that our i-Microglia can be co-cultured with iPSC-derived neurons, enabling studies of neuroimmune interactions. This is particularly relevant for modeling neurodegenerative and neuroinflammatory diseases such as Alzheimer’s disease, Parkinson’s disease, and multiple sclerosis.

In conclusion, our improved protocol provides a reproducible, xeno-free, and functionally validated system for generating human microglia from a widely adopted iPSC reference line, supporting a broad range of applications in neuroinflammation, disease modeling, and therapeutic screening.

## Acknowledgments

We thank Dr. Fernando Erra-Díaz for his suggestions regarding the phagocytosis and stimulation assays and Tina Beaulieu for administrative assistance. We also thank Dr. Deborah Kronenberg-Versteeg and Lena Erlebach from Hertie Institute for Clinical Brain Research, Tübingen, Germany for their assistance with microglia culture and immunofluorescence. This work was partially supported by the iPSC Neurodegenerative Disease Initiative (iNDI) funded to W.C.S (NIH 75N95020Q00265), and by JAX Scientific Service Process Improvement Funding to J.A.M. We gratefully acknowledge the contribution of the following Scientific Services at The Jackson Laboratory for expert assistance with the work described in this publication: Cellular Engineering, Flow Cytometry, Genome Technologies, Light Microscopy, and Scientific Instrument Services.

## Material and Methods

### iPSC Culture and Gene Editing

The KOLF2.1J human iPSC line was cultured under conditions described by Skarnes et al. (45). To generate the i-Microglia line, six transcription factors (TFs) were inserted into the CLYBL safe harbor locus using CRISPR/Cas9-mediated genome editing (46). The six TFs were divided into two sets of three, each driven by a TetON-inducible promoter and separated by T2A elements and integrated into the two CLYBL alleles (Figure 1).

Genome editing was performed as previously described (45). Cas9 ribonucleoprotein complexes were assembled using Alt-R S.p. HiFi Cas9 Nuclease V3 (IDT) and a synthetic guide RNA (Synthego) targeting CLYBL (CLYBL_sg1, Table 1). Two donor plasmids (J42 and J43, Table 1), each encoding a set of three TFs, were co-delivered.

**Table 1:**
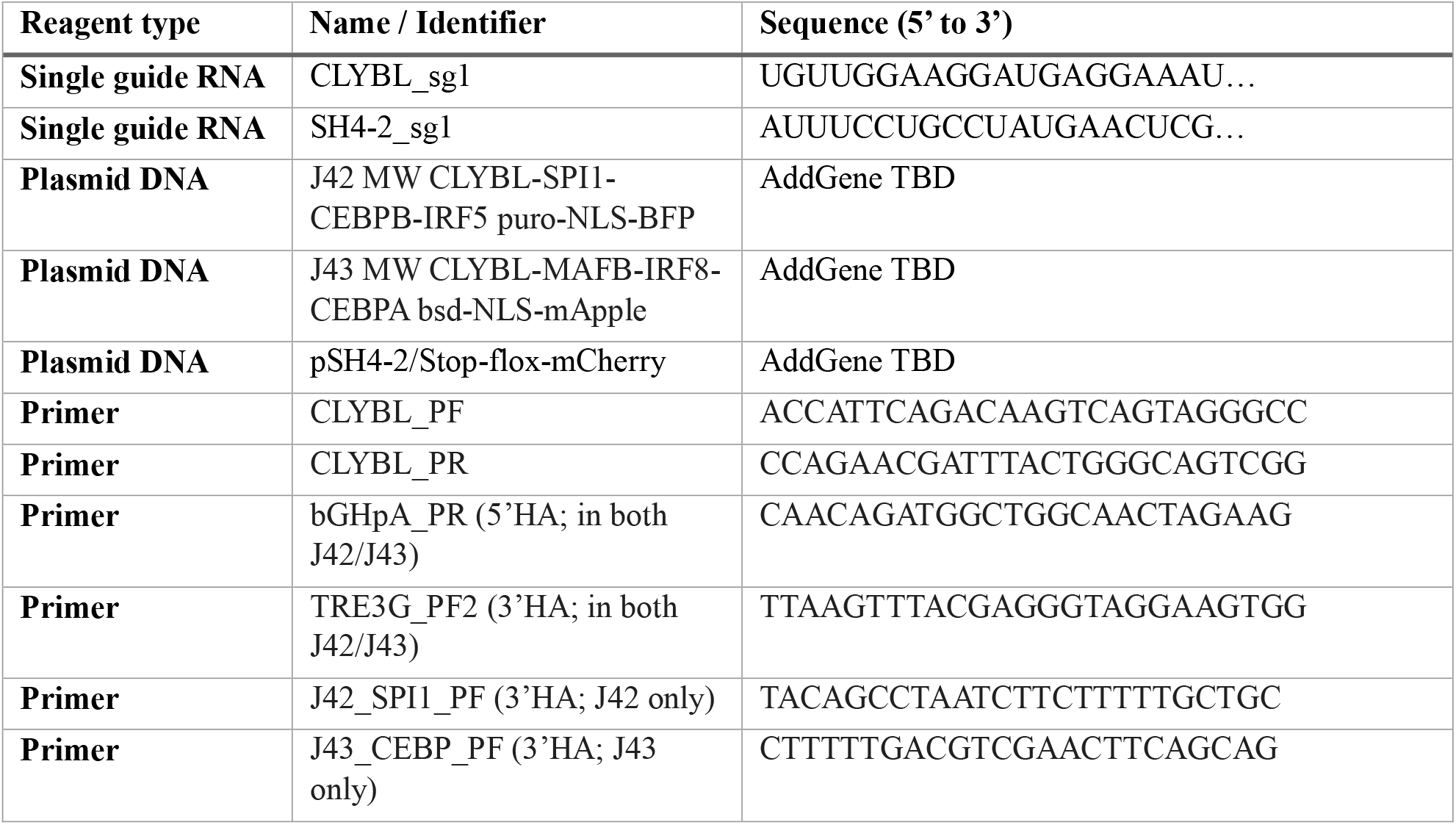
Reagents used for genome editing.

Briefly, 2 × 10□ Accutase-treated KOLF2.1J cells were pelleted and resuspended in 100 µL Primary Cell P3 buffer with supplement (Lonza), containing 20 µg Cas9 protein, 16 µg of sgRNA, and 2 µg of each donor plasmid. Plasmid DNA was pre-concentrated to ∼0.88 µg/µL in P3 buffer. Cells were nucleofected using the Amaxa 4D system (Lonza) and immediately recovered in StemFlex medium (Gibco) supplemented with RevitaCell (ThermoFisher) at 37°C.

One day post-nucleofection, the medium was changed to StemFlex alone and refreshed every other day. At day 3 selection was initiated with 40 µg/ml blasticidin and 0.5 µg/ml puromycin to enrich for cells with successful integration of both donors. Drug-resistant colonies were pooled, passed twice, and analyzed by flow cytometry to estimate the proportion of BFP/mApple-positive cells (∼80%).

To excise the selection cassettes, pooled cells were treated with 120U/ml of TAT-Cre recombinase (Millipore, stock 10,000 U/ml) for 6 hours. Excision efficiency was assessed by flow cytometry. Cells were then single cell plated as described by Skarnes et al. (45). Replicate plates were treated with or without blasticidin/puromycin. Clones from drug-sensitive wells on the untreated plate were expanded and genotyped by PCR across the left and right homology arms. Clones positive for both homology arms and negative for wild-type CLYBL were identified. Transgene-specific primers for J42 and J43 are listed in Table 1.

Selected clones were expanded into replicate vials and subjected to genome-wide SNP microarray analysis and G-band karyotyping. Clones A02 and D01, both exhibiting a normal male karyotype, were selected for further use.

### mCherry knock-in at SH4-2 safe harbor

To enable live-cell tracking, an mCherry reporter was integrated into the SH4-2 genomic safe harbor locus (20) in the A02 i-Microglia clone. Editing was performed using the same culture and nucleofection conditions described above, with a guide RNA targeting SH4-2 (SH4-2_sg1, Table 1) and SH4-2 donor plasmid encoding a CAG promoter, a loxP-flanked STOP cassette, and mCherry (pSH4-2/Stop-flox-mCherry, Table 1). To enhance and stabilize transgene expression, the construct included WPRE (Woodchuck Hepatitis Virus Posttranscriptional Regulatory Element) sequences, which improve transcript stability and translation efficiency (20).

Two days after replating, cells were treated with TAT-Cre for 6 hours to excise the STOP cassette and activate mCherry expression. Three days post-treatment, mCherry expression was assessed by flow cytometry in the bulk population and during clone isolation.

To enhance and stabilize transgene expression, the construct included WPRE (Woodchuck Hepatitis Virus Posttranscriptional Regulatory Element) sequences, which improve transcript stability and translation efficiency (20). Positive clones were expanded and re-evaluated after two passages to confirm stable mCherry expression.

### i-Microglia differentiation

Differentiation of 6-TF iPSC knockin clones into i-Microglia was performed with minor modifications to the protocol described by Dräger et al. (1). hiPSCs were maintained in mTeSR+ medium (StemCell Technologies) for at least two passages prior to differentiation. On day 0, cells were plated onto double-coated culture plates. The first coating consisted of Poly-D-Lysine (0.1 mg/mL, Gibco) prepared in Borate buffer (20X, ThermoFisher) diluted in culture-grade water (Gibco) and incubated at 37°C for 3 hours to overnight. This was followed by a second coating with Matrigel (Corning) diluted to 1x in DMEM/F12 (Gibco), incubated at 37°C for 2 hours.

For some experiments, a double coating of Poly-D-Lysine and Laminin 111 or Laminin-521 (Biolamina) was used. After Poly-D-Lysine coating as described above, plates were coated with 5 µg/ml of Laminin 111 or 521 for 18 hours at 4°C (for glass plates used in imaging) or for 2 hours at 37°C. Cells were seeded at the following densities: 27,000 cells per well in 24-well plates, 54,000 cells per well in 12-well plates, and 135,000 cells per well in six-well plates. For Laminin 111-coated plates, the seeding density was increased to 205,000 cells per six-well plate, with proportional adjustments for other formats. For confocal imaging, glass plates (Corning or CellView Advanced, Greiner Bio-One) were used and coated as described above.

Cells were seeded in mTeSR+ medium supplemented with 1x RevitaCell (ThermoFisher) and 2 µg/ml doxycycline (Sigma-Aldrich). On day 2, the base medium was replaced with Advanced DMEM/F12 (ThermoFisher) supplemented with 1% GlutaMAX (ThermoFisher), 100 ng/ml IL-34 (BioLegend), and 10 ng/ml GM-CSF (BioLegend). On day 4, a full medium change was performed, and 50 ng/ml of M-CSF (BioLegend) and 50 ng/ml of TGF-β1 (PeproTech) were added to the culture. On day eight, CX3CL1 (PeproTech) was added to the culture and doxycycline was withdrawn. From day 8 onward, the medium was changed every 2-3 days.

### Immunofluorescence

Cells were fixed at various time points using 4% formalin (Sigma-Aldrich) for 10 minutes at room temperature. Following fixation, cells were washed three times with 1x PBS for 5 minutes each, then permeabilized with 0.3% Triton X-100 (Sigma-Aldrich) for 15 minutes at room temperature. After an additional three PBS washes, cells were blocked with 5% bovine serum albumin (BSA; Sigma-Aldrich) for 1 hour at room temperature.

Primary antibody incubation was performed overnight at 4°C. The antibodies and their respective dilutions are listed in Table 2. Briefly, OCT4 and SSEA4 were used to characterize iPSCs, while IBA1, CD68, and PU.1 were used to assess i-Microglia identity.

**Table 2:** List of antibodies utilized in this study.

| Antibody | Vendor | Dilution | Application |
| --- | --- | --- | --- |
| Alexa 488 | ThermoFisher | 1:500 | IF |
| Alexa 568 | ThermoFisher | 1:500 | IF |
| CD68 | Cell Signaling | 1:150 | IF |
| IBA1 | Fuji Film | 1:300 | IF |
| **OCT4** | Cell Signaling | 1:300 | IF |
| **P2RY12** | Biolegend | 1:450 | FC |
| **PU.1** | ThermoFisher | 1:150 | IF |
| **SSEA4** | Cell Signaling | 1:300 | IF |
| **SSEA4** | Biolegend | 1:500 | FC |
| **DAPI** | ThermoFisher | 1:5000 | FC |
| **Hoechst** | Invitrogen | 2 µg/ml | Live Imaging |
| **CXCR1** | Biolegend | 1:400 | FC |
| **HLADR** | ThermoFisher | 1:1000 | FC |
| **CD40** | BD | 1:1000 | FC |
| **CD80** | BD | 1:1000 | FC |
| **CD83** | Biolegend | 1:500 | FC |
| **CD86** | Biolegend | 1:1000 | FC |
| **TUBB3 (TUJ1)** | Biolegend | 1:200 | IF |

Following primary antibody incubation, cells were washed three times with 1x PBS and incubated with Alexa Fluor 488- or Alexa Fluor 568-conjugated secondary antibodies for 1 hour at room temperature. After three additional washes, nuclei were stained with DAPI for 10 minutes. Samples were washed three final times with PBS before imaging.

Fluorescence imaging was performed using a Nikon Ti Eclipse Widefield microscope. Image processing and analysis were conducted using FIJI software (https://fiji.sc/).

### Phagocytosis assay

Phagocytic activity of i-Microglia was assessed using a modified version of the protocol described by Erra-Diaz et al. (47). On day 9 of differentiation, i-Microglia were incubated with 100 µg/mL of pHrodo^™^ Green *E. coli* BioParticles® (ThermoFisher, Cat# P35366) for 2 hours and 30 minutes at 37°C. Unloaded cells served as negative controls. To assess nonspecific particle binding, parallel samples were incubated with BioParticles under identical conditions at 4°C.

Following incubation, cells were harvested and analyzed by flow cytometry using a BD LSRFortessa^™^ cytometer. Phagocytic activity was quantified based on the fluorescence intensity of internalized particles.

For live-cell imaging and time-lapse analysis, i-Microglia were incubated with pHrodo^™^ Green BioParticles as described above. For time-lapse acquisition, cells were placed in an Incucyte® (Sartorius) live-cell imaging system, and images were captured every 15 minutes over a 4-hour period.

For high-resolution confocal imaging, cells were incubated with BioParticles for 1 hour, followed by nuclear staining with 2 µg/mL Hoechst 33342 (Invitrogen) for 20 minutes at 37°C. After staining, the medium was removed, cells were washed once with fresh medium and then replenished with fresh medium prior to imaging. Confocal images were acquired using a Dragonfly spinning disk confocal microscope (Andor).

### Flow cytometry analysis

Human iPSCs were dissociated as described by Skarnes et al. (45). Following dissociation, cells were washed once with 1x PBS to remove residual media and incubated with anti-SSEA4 antibody (Table 2) in FACS buffer (1x PBS + 2% fetal bovine serum, Gibco) for 30 minutes on ice. After incubation, cells were washed with PBS, centrifuged at 300 x g for 3 minutes, and resuspended in FACS buffer for analysis. Unstained cells were used as negative controls. Flow cytometry was performed using a BD LSRFortessa^™^ cytometer.

For i-Microglia, culture medium was aspirated, and cells were washed once with 1x PBS. Cells were then incubated with TrypLE Express (ThermoFisher) for 7–10 minutes at 37°C to facilitate detachment. Advanced DMEM/F12 medium was added to collect the cells, which were then centrifuged at 400 x g for 5 minutes. The pellet was resuspended in PBS, centrifuged again, and finally resuspended in FACS buffer with or without specific antibodies (see Supplementary Table 3). Cells were incubated on ice for 30 minutes.

For microglial identity, antibodies against P2RY12 and CX3CR1 were used. To assess immune activation, cells were stained with antibodies against HLA-DR, CD40, CD80, CD83, and CD86. Unstained and single-stained controls were included for gating and compensation.

Samples were analyzed on a BD LSRFortessa^™^ cytometer, and data were processed using FlowJo software (BD Biosciences).

### Stimulation of i-Microglia

i-Microglia differentiated for at least 10 days were stimulated with 500 ng/mL lipopolysaccharide (LPS; Sigma-Aldrich) for 22 hours. Media-only treated cells served as negative controls. Following stimulation, cells were harvested and processed for flow cytometry as described above.

### Co-culture of inducible cortical glutamatergic neurons with i-Microglia

Co-culture experiments were performed using iPSC-derived cortical glutamatergic neurons engineered to express NGN2 from the AAVS1 safe harbor locus, as described by Flores et al. (21). Neurons were differentiated for 11 days under the conditions established in that protocol.

On day 11, i-Microglia differentiated for 4 days on Laminin-521 (BioLamina) were dissociated using TrypLE Express (ThermoFisher) as previously described. To promote microglial attachment, 5 µg/mL Laminin-521 or 10 µg/mL Laminin-3D (R&D Systems) was added to the i-Microglia medium. A total of 0.35 × 10□ i-Microglia were seeded per well of a 24-well plate, each containing 0.35 × 10□ i-Neurons, maintaining a 1:1 cell ratio.

The medium in each well was also maintained at a 1:1 ratio of neuron and microglia media. Proliferation inhibitors (uridine and FDU), present in the original neuronal differentiation protocol, were withdrawn at the time of co-culture initiation. Cells were co-cultured for at least 10 days.

For immunofluorescence, cells were stained with βIII-tubulin (TUJ1) to label neurons and IBA1 to label microglia, following the protocol described by Marianita Santiana (Personal communication). Imaging was performed using a Nikon Ti Eclipse Widefield microscope, and images were processed using FIJI software (https://fiji.sc/).

### Flow cytometry data analysis

Flow cytometry data for hiPSC and i-Microglia characterization, as well as for quantification of mean fluorescence intensity (MFI), were analyzed using FlowJo software (version 10.10, BD Biosciences). Gating strategies were based on forward and side scatter profiles, exclusion of doublets, and fluorescence intensity relative to unstained controls.

### Quantification and statistical analysis

Quantitative data from flow cytometry experiments, including the number of biological replicates, mean values, and standard error of the mean (SEM), are provided in the corresponding figure legends. MFI values were calculated using FlowJo v10.10 and plotted using GraphPad Prism v10. Statistical significance was determined using unpaired two-tailed Student’s *t*-tests, with *p* < 0.05 considered significant.

