## Supplemental figures for "A xeno-free protocol for rapid differentiation of human iPSC-derived microglia from the KOLF2.1J reference line"

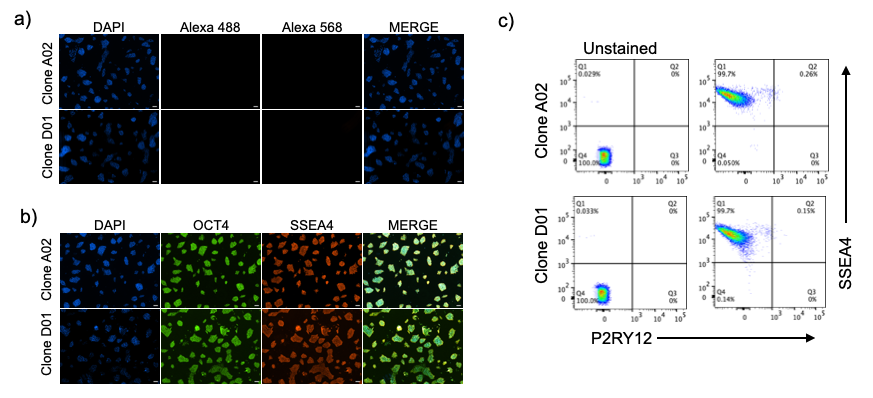

**Supplemental Figure S1.** Characterization of iMicroglia at the iPSC stage. A) Immunofluorescence (IF) for iPSC marker negative controls. B) Cells stained with OCT4 and SSEA4. C) Flow cytometry (FC) characterization using the iPSC marker SSEA4 and the microglia marker P2RY12.

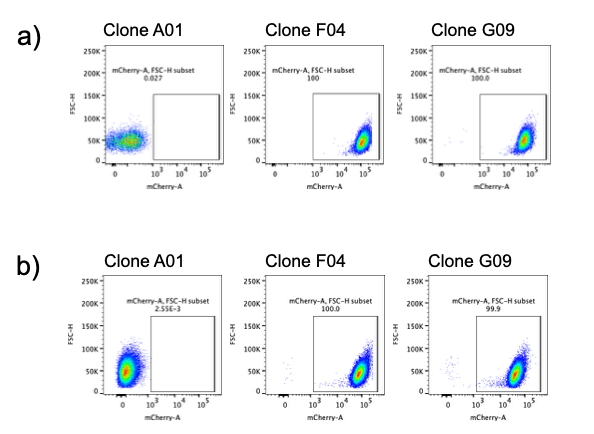

**Supplemental Figure S2.** Generation and validation of mCherry-expressing hiPSCs. (a) Flow cytometry analysis confirmed mCherry expression in SH4-2 knock-in clones F04 and G09. (b) Clones were expanded for two additional passages to verify stable mCherry expression. Clone A01, which lacked mCherry expression, was expanded as a negative control.

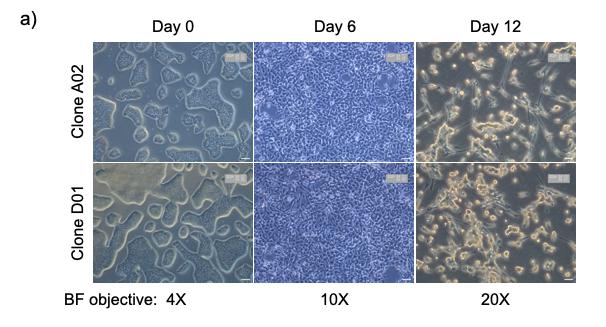

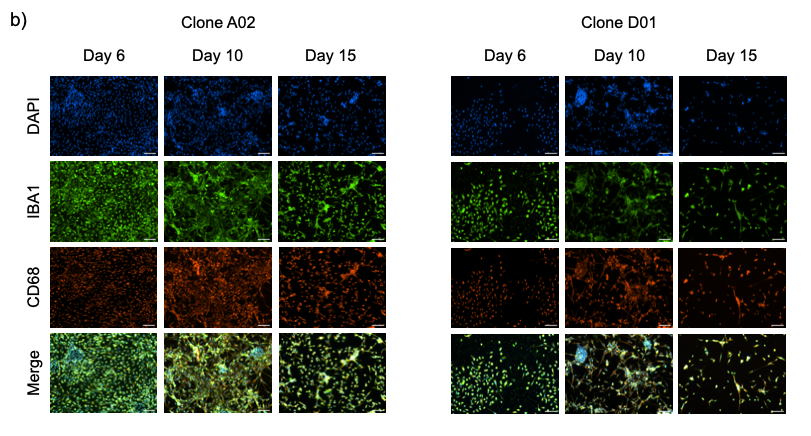

**Supplemental Figure S3**. Differentiation of i-Microglia on Poly-D-Lysine + Matrigel Substrate. (a) Brightfield (BF) images and (b) immunofluorescence (10× objective) showing the progression from undifferentiated iPSCs at day 0 to day 15 cells exhibiting mature microglial morphology and expression of microglial markers IBA1 and CD68. Data shown for KOLF2.1J i-Microglia clones A02 and D01.

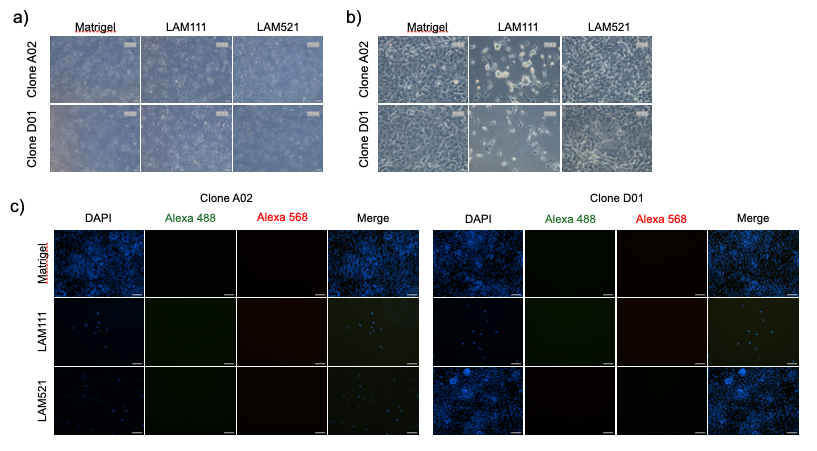

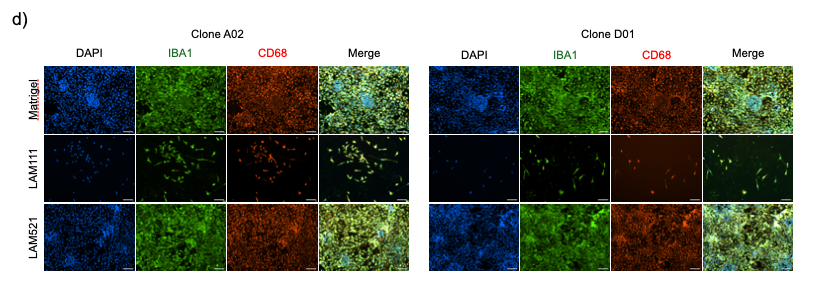

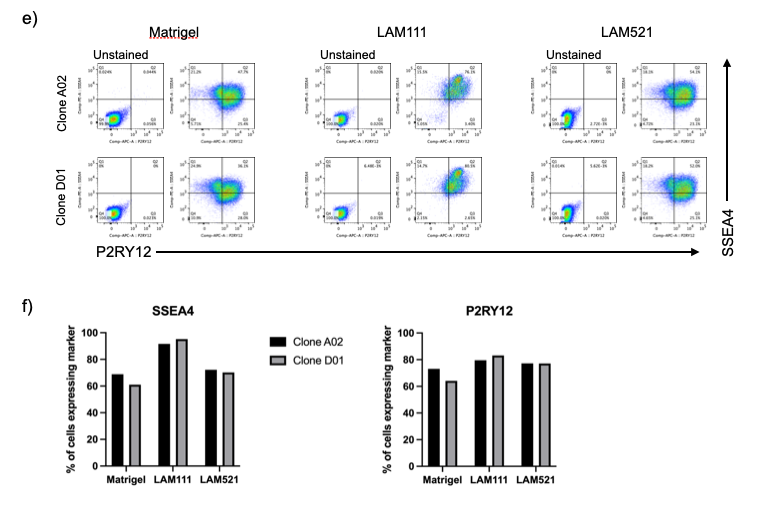

**Supplemental Figure S4**. Comparison of Substrate Conditions for i-Microglia Differentiation.(A–B) Representative brightfield images comparing differentiation on Matrigel, Laminin-111 (LAM111), and Laminin-521 (LAM521) at day 2 (A, 10× objective) and day 6 (B, 20× objective) for clones A02 and D01 (C) Negative control immunofluorescence (secondary antibody only) at day 6. (D) Immunofluorescence analysis at day [insert day] showing expression of IBA1 (green), CD68 (red), and DAPI (blue) in clones A02 and D01 across all three substrates. (E) Representative flow cytometry plots at day 6 showing SSEA4 (y-axis) and P2RY12 (x-axis) expression for each substrate condition in both clones. (F) Quantification of flow cytometry data showing the percentage of cells expressing SSEA4 (left) and P2RY12 (right) for clones A02 and D01 across Matrigel, LAM111, and LAM521 conditions.
